# Identification of anti-severe acute respiratory syndrome-related coronavirus 2 (SARS-CoV-2) oxysterol derivatives in vitro

**DOI:** 10.1101/2021.01.31.429001

**Authors:** Hirofumi Ohashi, Feng Wang, Frank Stappenbeck, Kana Tsuchimoto, Chisa Kobayashi, Wakana Saso, Michiyo Kataoka, Kouji Kuramochi, Masamichi Muramatsu, Tadaki Suzuki, Camille Sureau, Makoto Takeda, Takaji Wakita, Farhad Parhami, Koichi Watashi

**Affiliations:** Department of Virology II, National Institute of Infectious Diseases, Tokyo 162-8640, Japan; Department of Applied Biological Sciences, Tokyo University of Science, Noda 278-8510, Japan; MAX BioPharma, Inc., 2870 Colorado Avenue, Santa Monica, CA 90404, USA; The Institute of Medical Science, The University of Tokyo, Tokyo 108-8639, Japan; AIDS Research Center, National Institute of Infectious Diseases, Tokyo 162-8640, Japan; Department of Pathology, National Institute of Infectious Diseases, Tokyo 162-8640, Japan; Laboratoire de Virologie Moléculaire, Institut National de la Transfusion Sanguine, Paris 75739, France; Department of Virology III, National Institute of Infectious Diseases, Tokyo 208-0011, Japan; Institute for Frontier Life and Medical Sciences, Kyoto University, Kyoto 606-8507, Japan; MIRAI, JST, Saitama 332-0012, Japan

**Keywords:** SARS-CoV-2, COVID-19, oxysterols, antiviral, coronavirus, double membrane vesicle, replication, pharmacokinetics

## Abstract

Development of effective antiviral drugs targeting the severe acute respiratory syndrome-related coronavirus 2 (SARS-CoV-2) are urgently needed to combat the coronavirus disease 2019 (COVID-19). Oxysterols, defined as oxidized derivatives of cholesterol, include endogenous (naturally occurring) cholesterol metabolites as well as semi-synthetic oxysterol derivatives. We have previously studied the use of semi-synthetic oxysterol derivatives as drug candidates for inhibition of cancer, fibrosis, and bone regeneration. In this study, we have screened a panel of naturally occurring and semi-synthetic oxysterol derivatives for anti-SARS-CoV-2 activity, using a cell culture infection assay. We show that the natural oxysterols, 7-ketocholesterol, 22(*R*)-hydroxycholesterol, 24(*S*)-hydroxycholesterol, and 27-hydroxycholesterol, substantially inhibited SARS-CoV-2 propagation in cultured cells. Among semi-synthetic oxysterols, Oxy186 displayed antiviral activity comparable to natural oxysterols. In addition, related oxysterol analogues Oxy210 and Oxy232 displayed more robust anti-SARS-CoV-2 activities, reducing viral replication more than 90% at 10 μM and 99% at 15 μM, respectively. When orally administered in mice, peak plasma concentrations of Oxy210 fall into a therapeutically relevant range (19 μM), based on the dose-dependent curve for antiviral activity in our cell culture infection assay. Mechanistic studies suggest that Oxy210 reduced replication of SARS-CoV-2 with disrupting the formation of double membrane vesicles (DMVs), intracellular membrane compartments associated with viral replication. Oxy210 also inhibited the replication of hepatitis C virus, another RNA virus whose replication is associated with DMVs, but not the replication of the DMV-independent hepatitis D virus. Our study warrants further evaluation of Oxy210 and Oxy232 as a safe and reliable oral medication, which could help protect vulnerable populations with increased risk developing COVID-19.

## 1. Introduction

Coronavirus disease 2019 (COVID-19), caused by infection with the severe acute respiratory syndrome-related coronavirus 2 (SARS-CoV-2), has drastically impacted public health and, on a global scale, caused enormous harm to human societies and their economic vitality. In the search for effective treatments for COVID-19, understandably, the repurposing of existing FDA-approved drugs has been given high priority due to their known safety profiles [1]. For example, remdesivir (RDV), which was originally designed as an anti-ebola virus agent, has been repurposed to become the first and, to date, the only FDA-approved drug treatment for SARS-CoV-2 infection. Similarly, chloroquine and hydroxychloroquine, which are used to control malaria, have been investigated as COVID-19 treatments [2]. Beyond drug repurposing [3], other approaches are urgently needed to invigorate discovery research for new, specific, and potent anti-COVID-19 drugs.

Naturally occurring oxysterols include metabolites of cholesterol involved in the biosynthesis of steroid hormones, vitamin D, bile acids and other crucial signaling molecules [4, 5]. Beyond their role as passive and transient metabolites, endogenous oxysterols are increasingly recognized as lipid signaling molecules that can regulate a range of physiological processes, including lipid homeostasis, transport and metabolism as well as immune response [5]. In recent years, numerous reports have ascribed broad spectrum antiviral properties to naturally occurring oxysterols. For example, 20(*S*)-hydroxycholesterol [20(*S*)-OHC] and 22(*S*)-hydroxycholesterol [22(*S*)-OHC] reduced the infection of hepatitis B virus [6]; 25-hydroxycholesterol (25-OHC) and 27-hydroxycholesterol (27-OHC) displayed antiviral activities against herpes simplex virus [7], human papillomavirus-16, human rhinovirus [8], murine norovirus [9], rotavirus [10], and zika virus [11].

In this study, we focused on oxysterols, including naturally occurring and semi-synthetic oxysterols, to identify potent anti-SARS-CoV-2 agents, since we have already developed various semi-synthetic oxysterols as drug candidates in the context of cancer, fibrotic diseases and bone regeneration: Oxy133, an allosteric activator of Hedgehog (Hh) signaling, was designed for orthopedic applications, such as bone repair and spine fusion [12–14]; Oxy186, an inhibitor of Hh signaling that acts downstream of the Smoothened (Smo) receptor, was designed as a potential anti-tumorigenic agent [15], and Oxy210 was designed for application in cancer and fibrosis through dual inhibition of Hh and transforming growth factor-beta (TGF-β) signaling [16]. In the present report, using cell-based analysis, we demonstrate that Oxy210 and its analogue, Oxy232, display superior anti-SARS-CoV-2 activity compared to the natural oxysterols, 7-ketocholesterol (7-KC), 22(*R*)-hydroxycholesterol [22(*R*)-OHC], 24(*S*)-hydroxycholesterol [24(*S*)-OHC], and 27-OHC. Importantly, Oxy210 reduced viral replication and the formation of double membrane vesicles (DMVs), known RNA replication factories of coronaviruses and other RNA viruses [17–19]. Oral administration of a single dose of Oxy210 at 200 mg/kg in mice resulted in a peak plasma concentration (C_max_) of about 19 μM, which exceeds both the IC_50_ (5.6 μM) and IC_90_ (8.6 μM), respectively, determined in our cell-based assay. These data provide foundational evidence for Oxy210 and Oxy232 as potential anti-COVID-19 candidates for further therapeutic development in the future.

## 2. Results

### 2.1 Natural oxysterols have antiviral activity against SARS-CoV-2 infection

In this study, we used a cell-based SARS-CoV-2 infection system previously reported [20]. This infection system uses VeroE6 cells stably overexpressing the TMPRSS2 gene, which is a member of type II transmembrane serine proteases. Cells were treated with test compounds for 1 h during inoculation with a clinical isolate of SARS-CoV-2 at a multiplicity of infection (MOI) of 0.001, followed by washing out free virus and incubating the cells with test compounds for 24 h or 48 h (Figure. 1A and Materials and Methods). SARS-CoV-2 propagation in VeroE6/TMPRSS2 cells induced a cytopathic effect (CPE) at 48 h post-virus inoculation (Figure. 1B, panel b), and the treatment with remdesivir (RDV), a known replication inhibitor of SARS-CoV-2 [2], blocked the virus-induced CPE (Figure. 1B, panel c). SARS-CoV-2 propagation visualised by detecting viral nucleocapside (N) protein by immunofluorescence (IF) analysis was also blocked by RDV (Figure. 1C, panels b and c, red). Using this assay, we evaluated the antiviral effect of cholesterol and 7-ketocholesterol (7-KC) as a representative of natural oxysterols. 7-KC, but not cholesterol, reduced the SARS-CoV-2-induced CPE (Figure. 1B, panels d and e) and the spread of infection (Figure. 1C, panels d and e). To quantify antiviral activity, we measured viral RNA production in the culture supernatant, and cell viability upon treatment with natural oxysterols or cholesterol at 24 h post-inoculation. Cholesterol, 4beta-hydroxysterol (4beta-OHC) and 22(*S*)-hydroxycholesterol [22(*S*)-OHC], did not show apparent reductions in viral RNA, while 7-KC, 22(*R*)-hydroxycholesterol [22(*R*)-OHC], 24(*S*)-hydroxycholesterol [24(*S*)-OHC], and 27-hydroxysterol (27-OHC) reduced the production of viral RNA by 80-86% as compared to control (Figure 1D and supplementary Figure. S1). Evaluation of host cell viability showed no cytotoxic effect of the test compounds up to 30 μM, which is the maximum concentration in the SARS-CoV-2 infection assay shown in Figure. 1D, (Figure. 1E). These findings suggest that the oxysterols inhibited SARS-CoV-2 propagation without showing cytotoxicity.

**Figure 1.**
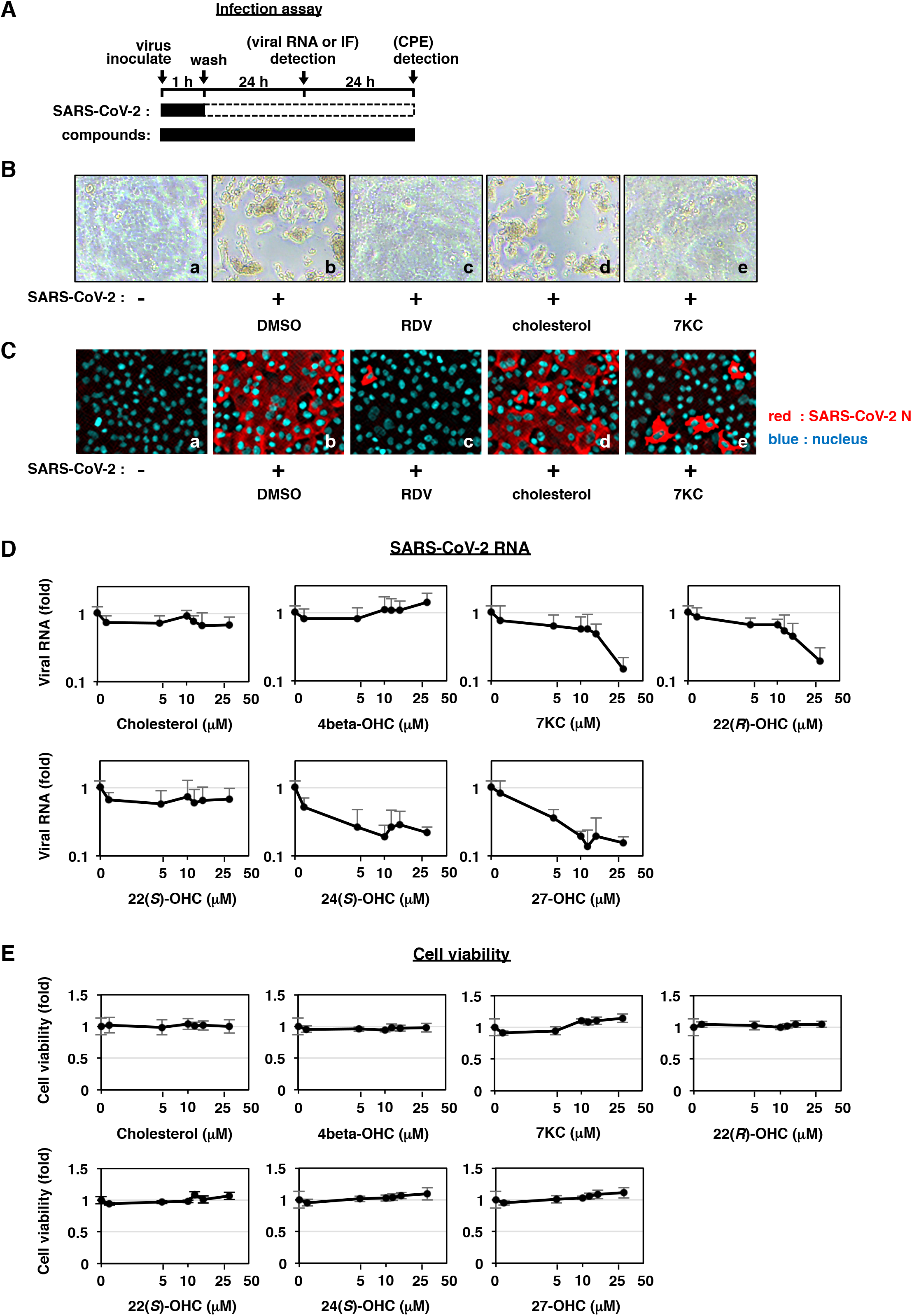
Oxysterols inhibit SARS-CoV-2 infection. **(A)** Schematic model of the SARS-CoV-2 infection assay. VeroE6/TMPRSS2 cells were inoculated with SARS-CoV-2 in the presence of compounds for 1 h, followed by washing out the free virus and incubating the cells with the compounds for 24 or 48 h. Viral RNA in the culture supernatant and viral N protein in the cells were quantified at 24 h post-inoculation by real time RT-PCR and immunofluorescence analyses, respectively. Cytopathic effects (CPE) were viewed under microscope at 48 h post-inoculation. Solid and dashed boxes indicate the periods that the cells were treated with and without the compounds or the virus, respectively. **(B)** Images of the cells treated with the virus in the presence of dimethyl sulfoxide (DMSO), 10 μM Remdesivir (RDV), 30 μM cholesterol, or 30 μM 7-ketocholesterol (7KC). **(C)** Viral N protein in the cells was detected by indirect immunofluorescence analysis. The red and blue signals represent viral N protein and nuclei, respectively. **(D)** Dose-response curves for SARS-CoV-2 RNA upon treatment with the compounds as indicated. Viral RNAs in the culture supernatant were quantified by real time RT-PCR and plotted against compound concentrations up to 30 μM. **(E)** Viability of cells treated with compounds as indicated for 24 h was quantified using MTT assay.

### 2.2 Semi-synthetic oxysterol derivatives, Oxy210, Oxy186 and Oxy232 inhibit the SARS-CoV-2 production

Although the natural oxysterols, 7-KC, 22(*R*)-OHC, 24(*S*)-OHC, and 27-OHC showed modest anti-SARS-CoV-2 activities, their physiological concentrations are at far below μM ranges [21, 22] in the circulation of healthy humans, suggesting their limited role, if any, in preventing viral infection in physiological condition. In search for oxysterols with improved antiviral activity, we evaluated the potential of semi-synthetic oxysterol derivatives for SARS-CoV-2 inhibition. SARS-CoV-2-induced CPE and virus propagation were blocked when treated with Oxy210 but not Oxy133 (Figure. 2A and 2B, panels d and e). Quantification of SARS-CoV-2 RNA in the culture supernatant at 24 h post-inoculation also showed that Oxy210 and its structurally related derivatives, Oxy186 and Oxy232, reduced viral RNA level in a dose-dependent manner, while Oxy133 did not show antiviral activity up to 15 μM (Figure 2C). The antiviral activity of Oxy186 was almost equivalent to that of the natural oxysterols shown earlier; the maximum reduction in viral RNA was 83% when used at 12 μM as compared to control (Figure 2C, note that the viral RNA shown in logarithm scale). On the other hand, Oxy210 and Oxy232 showed much higher antiviral potencies; viral RNA production was reduced by 99.4% (Oxy210) and 99.9% (Oxy232) at the maximum at 15 μM (Figure 2C). No significant cytotoxicity by Oxy186 and Oxy210 was found up to 15 μM, the maximum concentration in the infection assay, however, Oxy232 slightly reduced cell viability when used at concentrations above 10 μM (Figure 2D). Due to the greater availability of Oxy210 we performed further studies with this oxysterol analogue. The 50% and 90% maximal inhibitory concentration (IC_50_, IC_90_) and 50% maximal cytotoxic concentration (CC50) of Oxy210 were 5.6 μM, 8.6 μM, and >15 μM, respectively.

**Figure 2.**
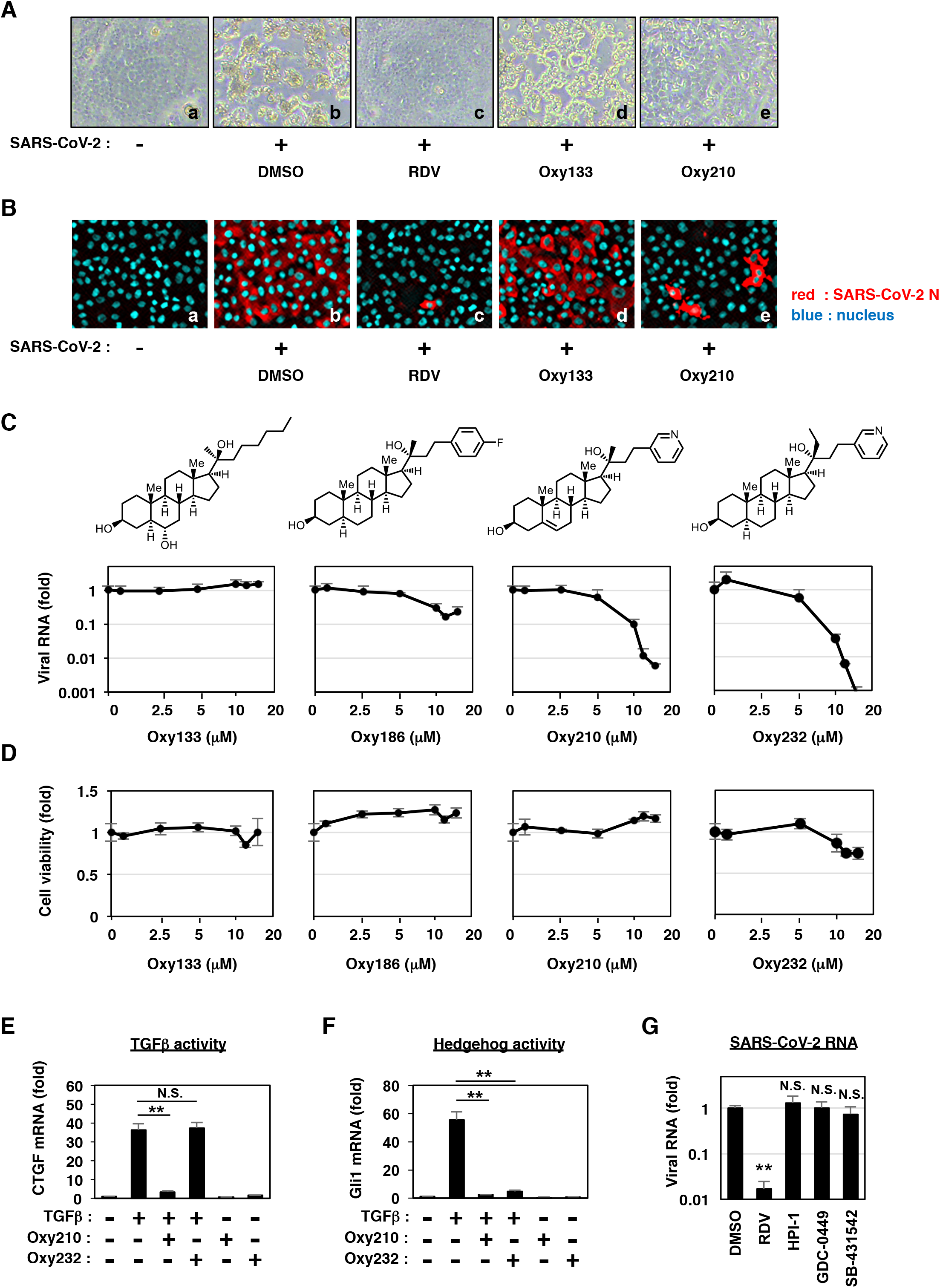
Oxy210, an oxysterol derivative, potently inhibits the SARS-CoV-2 propagation and alleviates the virus-induced CPE. **(A)** Virus-induced CPE was examined in the cells inoculated with the virus in the presence of DMSO, 10 μM RDV, 10 μM Oxy133, or 10 μM Oxy210. **(B)** Viral N protein in the cells was detected by immunofluorescence analysis as described in Figure 1C. **(C)** Dose-response curves for viral RNA upon treatment with oxysterol derivatives as indicated. The secreted viral RNA in the culture supernatant at 24 h post-inoculation was quantified by real time RT-PCR and plotted against compound concentration. The chemical structures of oxysterols are also shown above the graphs. **(D)** Viability of cells treated with the compounds was quantified using MTT assay. **(E, F)** Inhibitory effects toward TGFβ and Hh signaling. NIH3T3 cells pretreated with or without Oxy210 or Oxy232 for 2 h were stimulated with 20 ng/mL TGFβ (E) or with conditioned medium from CAPAN-1 human pancreatic tumor cells that contain Shh (F) [15, 16] in the presence or absence of the compounds. Cellular mRNAs were extracted to quantify a TGFβ-downstream gene, connective tissue growth factor (CTGF) (E), a Hh target gene, Gli1 (F), and Oaz1 for normalization of CTGF and Gli1 (E, F). **(G)** At 24 h post inoculation, Viral RNA produced from the cells treated with DMSO, 10 μM RDV, 10 μM HPI-1, 10 μM GDC-0449, or 10 μM SB-431542, was quantified with real time RT-PCR. All data are shown with error bars indicating S.D, ** P < 0.01 vs. DMSO; N.S., not significant, with Student’s t-test.

We previously reported that Oxy210 inhbited Hedgehog (Hh) and transforming growth factor β (TGFβ) signalings in fibroblastic cells and tumor cells [16]. In contrast, Oxy232, a close structural analogue of Oxy210, is devoid of significant TGFβ inhibitory properties (Figure 2E), but retains inhibitory activity toward Hh signaling (Figure 2F), suggesting that inhibition of TGFβ signaling is not resposible for the anti-SARS-CoV-2 activity. Consistent with this observation, treatment with the TGFβ signaling inhibitor, SB431542, did not significantly inhibit the production of viral RNA (Figure 2G). In addition, inactivation of Hh pathway by either HPI-1 or GDC0449 did not decrease the viral RNA levels (Figure 2G). These data suggest that Oxy210, Oxy232 and other antiviral oxysterol analogues inhibit SARS-CoV-2 production independent of the inhibition of Hh or TGFβ signaling pathways.

### 2.3 Oxy210 inhibits the intracellular SARS-CoV-2 replication and formation of double membrane vesicles

To determine which steps in the SARS-CoV-2 life cycle (Figure. 3A, left) were inhibited by Oxy210, we performed a time of addition assay (Figure. 3A, upper right). We examined the antiviral effect of Oxy210 in three different experimental groups, with different compound treatment times (Figure. 3A, a-c); (a) Compounds were treated during the 1 h virus inoculation and the additional 23 h up to detection to represent the whole life cycle (a, blue); (b) Compounds were present during the 1 h virus inoculation and an additional 2 h, and then removed to represent the virus entry process (b, green); and (c) Compounds were added 2 h after virus inoculation and were present for the remaining 21 h to represent the post-entry period (c, orange). We confirmed that Chloroquine (CLQ), a reported SARS-CoV-2 entry inhibitor that acts through modulation of intracellular pH [2, 23, 24], showed the most inhibitory effect when introduced in the entry-step of infection (Figure 3A, lower right, lane 8). [Because of the multiple rounds of viral re-infection in our assay, entry inhibitors can also show antiviral effects when introduced at post-entry (Figure 3A, lower right, lane 9)]. We also confirmed the mode of action of RDV, a reported inhibitor of intracellular viral RNA replication [25], by showing no significant effect on the virus entry-step (Figure 3A, lower right, lane 5) and a remarkable inhibition of post-entry phase (Figure 3A, lower right, lane 6). In this assay system, Oxy210, but not the negative control Oxy133, clearly reduced viral RNA levels when present during the whole life cycle and the postentry, but not at the entry phase, similar to the effects of RDV (Figure 3A, lower right, lanes 10-12). This finding suggests that Oxy210 targets intracellular virus replication, rather than viral entry.

**Figure 3.**
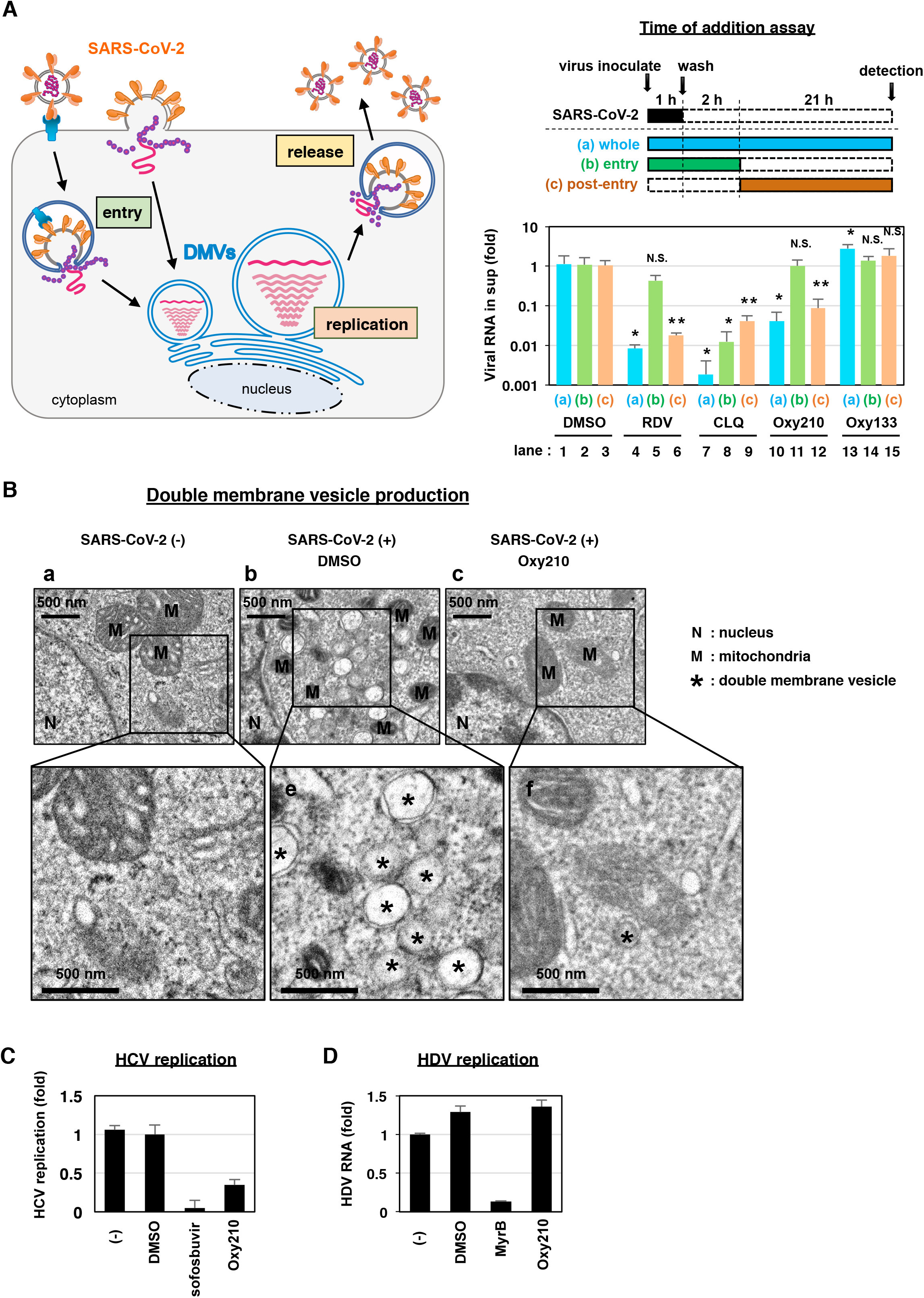
Oxy210 inhibits the SARS-CoV-2 genome replication. **(A)** Determination of the target step of compounds in the SARS-CoV-2 life cycle using time of addition analysis. The left diagram shows the life cycle of SARS-CoV-2, including the steps for viral entry, replication, and release. The upper right diagram shows the experimental procedures of the time of addition analysis. The assay was performed under three different conditions (a, whole; b, entry; and c, post-entry): (a) the cells were treated with the compounds for 24 h throughout the whole procedure (whole life cycle); (b) compounds were added during the 1 h virus inoculation and then removed after an additional 2 h treatment (entry); (c) compounds were added at 2 h postinoculation and presented for the remaining 21 h until harvest (post-entry). Solid and dashed boxes indicate the periods of presence and absence of the compounds, respectively. The lower right graph shows the real time RT-PCR quantified viral RNA produced from the cells treated with 15 μM RDV, 15 μM CLQ, 10 μM Oxy210 or 10 μM Oxy133 under the three experimental conditions. All data are shown with error bars indicating S.D, * P < 0.05 vs. DMSO; ** P < 0.01 vs. DMSO; N.S., not significant; with Student’s t-test. **(B)** SARS-CoV-2 infected (panel b and c) or uninfected (panel a) cells were treated with the compounds (b, DMSO; c, 10 μM Oxy210) as indicated and examined with electron microscopy. Images in panels d, e, and f show the insets in panels a, b, and c, respectively, at higher magnification. N, nucleus; M, mitochondria; *, double-membrane vesicle. **(C)** Hepatitis C virus (HCV) replication was evaluated by measuring the luciferase activity in LucNeo#2 cells carrying the discistronic HCV NN (genotype-1b) subgenomic replicon RNA and the luciferase gene (see Materials and Methods), treated with or without DMSO, 10 μM Oxy210, or 1 μM sofosbuvir as a positive control for 48 h. **(D)** Hepatitis D virus (HDV) replication was measured by quantifying HDV RNA using real time RT-PCR in HepG2-hNTCP-C4 cells infected with HDV and treated with or without DMSO or 10 μM Oxy210 for 6 days. 200 nM myrcludex-B (MyrB) was used as a positive control to inhibit HDV infection.

Coronaviruses generally induce the formation of unique membrane compartments, called double membrane vesicles (DMVs), which enables an efficient viral RNA replication [18, 26]. We found that DMV formation occurs with infection by SARS-CoV-2 in VeroE6/TMPRSS2 cells (Figure. 3B, panels b and e, *). Interestingly, treatment with Oxy210 remarkably reduced the DMV formation in the SARS-CoV-2-infected cells (Figure. 3B, panels c and f, *). We examined the speficicity of Oxy210’s effect on DMV dependent virus replication by evaluating the antiviral effect on hepatitis C virus (HCV) and hepatitis D virus (HDV), which are other RNA viruses that drive replication in a DMV-dependent and -independent manner, respectively [26, 27]. Similar to the effect of an HCV polymerase inhibitor, sofosbuvir, used as a positive control, Oxy210 reduced the DMV-dependent RNA replication of HCV (Figure 3C), while the antiviral activity was not observed in HDV infection that was inhibited by the positive control, MyrB (Figure 3D). These data are consistent with the idea that Oxy210 specifically inhibits the DMV-dependent virus replication, although it remains to be determined whether Oxy210 directly inhibits the DMV formation machinery, which we will examine in future studies (see the discussion below).

### 2.4 Pharmacokinetics of Oxy210 in mice

Given its higher anti-SARS-CoV-2 potency compared to the natural oxysterols, we questioned whether oral administration of Oxy210 in mice would result in plasma concentrations high enough to sustain significant antiviral activity *in vivo.* According to a previously established protocol [16], a single dose of Oxy210 at 200 mg/kg was orally administered to mice and the plasma concentration examined at 0.25, 0.5, 1, 2, 4, and 8 hours (h). Oxy210 was well tolerated by the mice in this experiment. After 1 h (T_max_), Oxy210 reached a peak plasma concentration (C_max_) of 8,155 ng/mL (19.4 μM) with an overall exposure of 29,305 h*ng/mL, as measured by the area under the curve (AUC) (Figure. 4)

**Figure 4.**
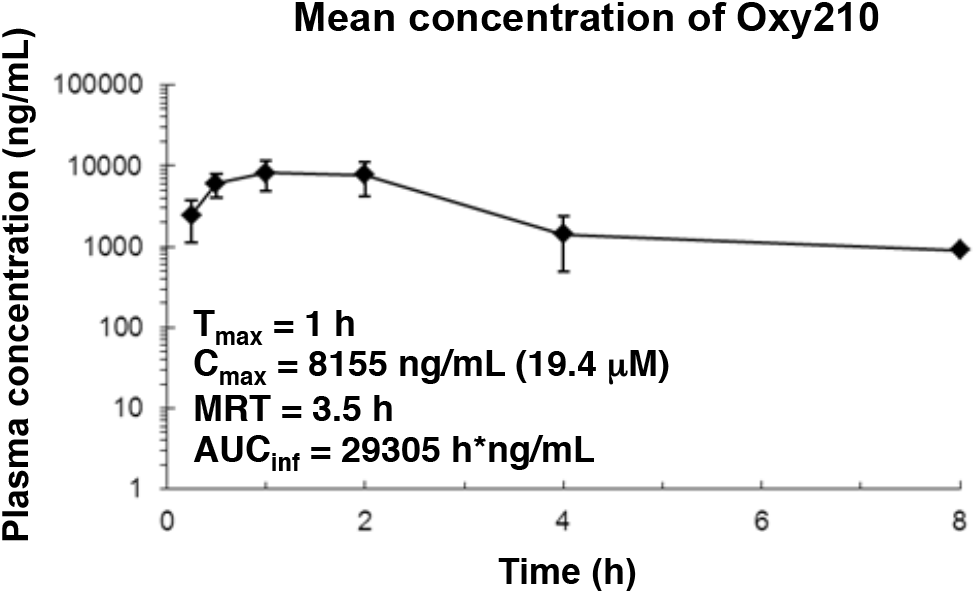
Pharmacokinetics of Oxy210 in mice. A single dose of Oxy210 at 200 mg/kg, formulated in 3% DMSO + 7% Ethanol + 5% PEG400 + 85% corn oil, was administered to balb/c mice by oral gavage. Plasma samples were taken at 0.25, 0.5, 1, 2, 4, and 8 h, followed by LC/MS analysis of the plasma to quantify Oxy210 concentrations.

In a separate study, Oxy210 was administered to mice via a chow diet containing 4 mg Oxy210/g of food. Oxy210 plasma concentrations were measured at 24, 48, and 96 h. No adverse effects or significant loss of body weight was recorded during this 96 h experiment. Accumulation of Oxy210 in plasma was greatest after 96 h, averaging at 2,682 ng/mL (6.4 μM) (Supplementary Table. S1). The concentration in the liver and the lung after 96 h was higher at 6,869 ng/mL (16.3 μM) and 4,137 ng/mL (9.8 μM), respectively (Supplementary Table. S1). These pharmacokinetics data indicate that oral administration of Oxy210 in mice, via oral gavage or mixed into a chow diet, results in plasma and lung concentrations that could sustain signifcant anti-SARS-CoV-2 activity *in vivo*.

## 3. Discussion

In this study, we have evaluated the anti-SARS-CoV-2 activity of a collection of naturally occurring and semi-synthetic oxysterol derivatives in cell cultures. Oxysterols are a class of understudied molecules that, until recently, have rarely been considered as a source of therapeutic drug candidates. In fact, most naturally occurring oxysterols cannot be ideal drug candidates for several different reasons, such as metabolic instability and overlapping biological activities. For example, 25-OHC, in addition to its antiviral properties, can also amplify the activation of immune cells and increases the production of potentially harmful immune mediators which are linked to the development of atherosclerosis [5]. Semi-synthetic oxysterol derivatives, by contrast, often posess improved drug like properties, in terms of potency, selectivity, metabolic stability and drug safety characteristics, compared to their naturally occuring counterparts. Given the urgency of the COVID-19 pandemic, we seek to establish a suitable drug development candidate with potent anti-SARS-CoV-2 activity that does not elicit unrelated or untoward pharmacological responses. In this study, Oxy210, a semi-synthetic oxysterol, was identified as a potent replication inhibitor of the SARS-CoV-2 which reduced the formation of DMVs. The peak plasma concentration of Oxy210 reached after administration via oral gavage (19 μM) and the plasma (6.4 μM) and lung (9.8 μM) concentrations reached after administration through diet, fall into a therapeutically meaningful range as they approach or exceed the IC_50_ (5.6 μM) and IC_90_ (8.6 μM) concentrations observed in our cell based assays. Therefore, Oxy210 and its analogues, such as Oxy232, could potentially serve as drug candidates targeting COVID-19.

We have previously characterized Oxy210 as a Hh and TGFβ signaling inhibitor [16] and have demonstrated protective effects of Oxy210 in a mouse model of idiopathic pulmonary fibrosis (IPF) (Parhami et al., unpublished observations). Oxy232, a close structural analogue of Oxy210, but devoid of significant TGFβ pathway inhibitory properties, displayed anti-SARS-CoV-2 activity comparable to Oxy210, suggesting that the mechanisms of the anti-SARS-CoV-2 activity shared by Oxy210 and Oxy232 are likely unrelated to TGFβ inhibitory properties exhibited by Oxy210. This notion was further supported by the lack of antiviral activity displayed by a TGFβ pathway inhibitor, SB431542 (10 μM). Also, the antiviral activity is not likely to be due to the inhibition of Hh pathway, as suggested by the lack of antiviral activity of Hh pathway inhibitors, HPI-1 (10 μM) and GDC0449 (10 μM). Given the absence of unrelated biological activities, such as TGFβ inhibition, Oxy232 may be a preferable drug candidate compared to Oxy210.

A recent publication reported significant anti-SARS-CoV-2 activity of 27-OHC; low concentrations of 27-OHC inhibited post-entry and higher concentration inhibited viral entry process [21]. Our time-of-addition analysis suggests that Oxy210 predominantly inhibits the post-entry process, which includes viral RNA replication in the replication factory and the following assembly of progeny virus and its secretion. We observed the formation of DMVs in SARS-CoV-2-infected cells, as previously reported [18, 28]. DMVs, membrane compartments separated from the nuclease/protease-rich cytosol, are generally considered to be sites for efficient replication of genomic RNA of coronaviruses and of certain other RNA viruses, such as HCV [26]. DMVs are also very likely to be important in SARS-CoV-2 replication [17]. We showed that the production of DMVs, induced by SARS-CoV-2, was substantially reduced with Oxy210 treatment. Antiviral effects of Oxy210 were also observed during the replication of HCV, a virus that depends on DMVs for replication, but not with HDV, a virus that replicates independently of DMVs. These findings suggest that Oxy210 specifically reduces DMV-dependent virus replication. It is not clear, however, whether Oxy210 directly inhibits the formation of DMVs. Further studies will be performed in the future to analyze the mode of action for the antiviral activity of Oxy210 and its analogues, and to elucidate the molecular mechanisms underlying their inhibitory effects on SARS-CoV-2 replication and DMV formation.

It is noteworthy to outline the potential advantages of semi-synthetic oxysterols as anti-SARS-CoV-2 agents, compared to established antiviral compounds, such as RDV:

1. RDV has to be administered intravenously, most often in a hospital setting, whereas the oxysterols could potentially be dosed orally (via a pill or liquid gel). A safe and reliable oral medication could be administered at an earlier stage, at the time of confirming SARS-CoV-2 infection, and potentially benefit asymptomatic individuals and those at high risk of infection who have the close-contact with infected individuals and medical care workers as a prophylactic treatment.
2. Oxysterols reprogram the host cell, interfering with the ability of the virus to use its machinery to replicate, reducing the likelihood of emerging drug resistance and likely possess universal antiviral activity against SARS-CoV-2 mutant strains.
3. The scale up and manufacturing of oxysterol-based drug candidates is expected to be straightforward and process friendly, especially when compared to the manufacturing process of RDV, which is rather difficult to prepare on scale.

We conclude that semi-synthetic oxysterol derivatives, such as Oxy210 and Oxy232, could be promising leads in the search for COVID-19 drug candidates, used alone, or in combination with other therapies currently FDA approved or under investigation, such as RDV, convalescent plasma or antibody treatments.

## 4. Materials and Methods

### Compounds and the synthesis of oxysterol derivatives

Commercially available oxysterols were obtained from Sigma Aldrich. Oxy133, Oxy186 and Oxy210 were prepared as previously described [12, 15, 16]. Oxy232 was prepared via a similar three-step synthesis described for Oxy186 and Oxy210, except for using ethyl magnesium bromide (instead of methyl magnesium bromide or methyl lithium) in step three. RDV was purchased from Chemscene; CLQ was purchased from Tokyo Chemical Industry; GDC-0449 was purchased from APExBIO, HPI-1 and SB-431542 was purchased from Cayman Chemical, sofosbuvir was purchased from MedChemExpress, and Myrcludex-B was synthesized by Scrum.

### Cell culture

VeroE6/TMPRSS2 cells, VeroE6 cells overexpressing transmembrane protease, serine 2 (TMPRSS2) [20, 29], were cultured in Dulbecco’s modified Eagle’s medium (DMEM; Wako) supplemented with 10% fetal bovine serum (FBS; Cell Culture Bioscience), 10 units/mL penicillin, 10 μg/mL streptomycin, 10 mM HEPES (pH 7.4), and 1 mg/mL G418 (Nacalai) at 37°C in 5% CO2. During the infection assay, 10% FBS was replaced with 2% FBS and G418 removed. LucNeo#2 cells, carrying HCV subgenomic replicon, were kindly provided by Dr. Kunitada Shimotohno at National Center for Global Health and Medicine [30] and were cultured in DMEM supplemented with 10% FBS, 10 units/mL penicillin, 10 μg/mL streptomycin, 0.1 mM nonessential amino acids (Invitrogen), 1 mM sodium pyruvate, 10 mM HEPES (pH 7.4), and 0.5 mg/mL G418 at 37°C in 5% CO2. HepG2-hNTCP-C4 cells, a HepG2 cell clone overexpressing the HDV entry receptor, sodium taurocholate cotrasporting polypeptide (NTCP), and highly susceptible to HDV infection [6] were cultured in GlutaMax (Invitrogen) supplemented with 10 units/ml penicillin, 10 μg/ml streptomycin, 10% FBS, 10 mM HEPES (pH 7.4), 50 μM hydrocortisone, and 5 μg/ml insulin at 37°C in 5% CO2.

### SARS-CoV-2 infection assay

SARS-CoV-2 was handled in a biosafety level 3 (BSL3) facility. We used the SARS-CoV-2 Wk-521 strain, a clinical isolate from a COVID-19 patient, and obtained viral stocks by infecting VeroE6/TMPRSS2 cells [20]. VeroE6/TMPRSS2 cells were inoculated with SARS-CoV-2 at an MOI of 0.001 (Figure 1B, 1C, 2A, and 2B), 0.003 (Figure 1D, 2C, and 3A), and 1 (Figure 3B) for 1 h and unbound virus removed by washing. Cells were cultured for 24 h prior to measuring extracellular viral RNA or detecting viral encoded N protein, for 48 h to detect cytopathic effects (CPE), and for 7 h to observe cells by electron microscopy. Compounds were added during virus inoculation (1 h) and after washing (24 or 48 h) except for time of addition assay shown in Figure 3A.

For the time of addition assay, we added compounds with three different timings (Figure 3A): (a) present during the 1 h virus inoculation and maintained throughout the 23 h infection period (whole life cycle); (b) present during the 1 h virus inoculation and for an additional 2 h and then removed (entry); or (c) added at 2 h after virus inoculation and present for the remaining 21 h until harvest (post-entry). Inhibitors of viral replication such as remdesivir (RDV) are expected to show antiviral activity in (a) and (c), but not (b), while entry inhibitors including CLQ reduce viral RNA in all three conditions [2].

### Quantification of viral RNA

Viral RNA in the culture supernatant was extracted with a QIAamp Viral RNA mini (QIAGEN) or MagMax Viral/Pathogen II Nucleic Acid Isolation kit (Thermo Fisher Scientific) and quantified by real time RT-PCR analysis with a one-step qRT-PCR kit (THUNDERBIRD Probe One-step qRT-PCR kit, TOYOBO) using 5’-ACAGGTACGTTAATAGTTAATAGCGT-3’, 5’-ATATTGCAGCAGTACGCACACA-3’, and 5’-FAM-ACACTAGCCATCCTTACTGCGCTTCG-TAMRA-3’ (E-set) [31].

### Detection of viral N protein

Viral N protein was detected using a rabbit anti-SARS-CoV N antibody [32] as a primary antibody with AlexaFluor 568 anti-rabbit IgG or anti-rabbit IgG-HRP (Thermo Fisher) as secondary antibodies together with DAPI to stain the nucleus by indirect immunofluorescence as described previously [33].

### Quantification of cell viability

Cell viability was determined by MTT assay as previously reported [33].

### Quantification of transforming growth factor (TGF)-β and Hedgeog (Hh) activity

TGFβ activity was examined with NIH3T3 cells precultured with DMEM containing 0.1% bovine calf serum (BCS) overnight. NIH3T3 cells were pretreated with the compounds for 2 h and then stimulated with TGFβ1 (20 ng/mL) in the presence or absence of compounds. After 48 h, RNA was extracted from the cells and analyzed for quantifying the mRNA for a TGFβ target gene, connective tissue growth factor (CTGF), and Oaz1 for normalization. For examination of Hh activity, NIH3T3 cells pretreated with the compounds for 2 h were treated with conditioned medium from CAPAN-1 human pancreatic tumor cells that contain Shh in the absence or presence of the compounds. Cellular RNA was extracted and analyzed for the expression of a Hh target gene, Gli1, and normalized to Oaz1 expression.

### Electron microscopic analysis

Cell were fixed with the buffer [2.5% glutaraldehyde, 2% paraformaldehyde, and 0.1 M phosphate buffer (pH 7.4)] for 1 h at room temperature followed by with 1% osmium tetroxide, stained in 1% uranyl acetate, dehydrated through a graded series of alcohols and embedded in Epon. Ultrathin sections were stained with uranyl acetate and lead citrate to observe with a transmission electron microscope (HT7700; Hitachi, Ltd., Japan)

### Hepatitis C virus (HCV) replication assay

HCV replication activity was measured using LucNeo#2 cells, carrying a subgenomic replicon RNA for an HCV NN strain (genotype-1b) and the luciferase gene driven by the HCV replication [30]. LucNeo#2 cells were treated with the compounds indicated in Figure 3C for 48 h and the luciferase activity was measured with Luciferase Assay System kit (Promega). Sofosbuvir, a clinically used HCV polymerase inhibitor, was used as a positive control.

### Hepatitis D virus (HDV) replication assay

HDV were recovered from the culture supernatant of Huh7 cells transfected with the plasmids for HDV genome and for hepatitis B virus surface antigen [34]. HepG2-hNTCP-C4 cells were inoculated with HDV for 16 h and were further cultured for 6 days in the presence or absence of Oxy210 to detect intracelular HDV RNA [34]. Myrcludex-B (Myr-B), used as a positive control that inhibits HDV infection, was treated during the virus inoculation.**Pharmacokinetics of Oxy210 in mice.** We perfomed the pharmacokinetics analysis in mice by oral administration with Oxy210 as described previously [16].

## Author Contributions

Conceptualization, F.P., K.W.; investigation, H.O., F.W., F.S., K.T., C.K., W.S., M.K., K.W.; analysis, all authors; resources, T.S., C.S., M.T.; writing-original draft preparation, H.O., F.S., F.P., K.W.; supervision, F.P., K.W.; funding acquisition, H.O., T.S., M.T., F.P., K.W. All authors have read and agreed to the published version of the manuscript.

## Acknowledgments

The HCV subgenomic replicon, LucNeo#2 cells, was kindly provided from Dr. Kunitada Shimotohno at National Center for Global Health and Medicine. The HDV-encoding plasmid was a generous gift from Dr. John Taylor at the Fox Chase Cancer Center.

## Abbreviations

SARS-CoV-2: severe acute respiratory syndrome-related coronavirus 2
COVID-19: coronavirus disease 2019
PK: pharmacokinetics
DMVs: double membrane vesicles
RDV: remdesivir
CPE: cytopathic effect
N protein: nucleocapsid protein
7KC: 7-ketocholesterol
4beta-OHC: 4beta-hydroxycholesterol
22(*S*)-OHC: 22(*S*)-hydroxycholesterol
22(*R*)-OHC: 22(*R*)-hydroxycholesterol
24(*S*)-OHC: 24(*S*)-hydroxycholesterol
27-OHC: 27-hydroxycholesterol
IC_50_: 50% maximal inhibitory concentration
IC_90_: 90% maximal inhibitory concentration
CC50: 50% maximal cytotoxic concentration
CLQ: chloroquine
HCV: hepatitis C virus
HDV: hepatitis D virus
CTGF: connective tissue growth factor

## References

1. Watashi, K., Identifying and repurposing antiviral drugs against severe acute respiratory syndrome coronavirus 2 with in silico and in vitro approaches. Biochem Biophys Res Commun, 2020.

2. Wang, M., et al., Remdesivir and chloroquine effectively inhibit the recently emerged novel coronavirus (2019-nCoV) in vitro. Cell Research, 2020. 30(3): p. 269–271.

3. Edwards, A., What Are the Odds of Finding a COVID-19 Drug from a Lab Repurposing Screen? J Chem Inf Model, 2020. 60(12): p. 5727–5729.

4. Griffiths, W.J. and Y. Wang, Oxysterol research: a brief review. Biochemical Society Transactions, 2019. 47(2): p. 517–526.

5. Luchetti, F., et al., Endothelial cells, endoplasmic reticulum stress and oxysterols. Redox Biology, 2017. 13: p. 581–587.

6. Iwamoto, M., et al., Evaluation and identification of hepatitis B virus entry inhibitors using HepG2 cells overexpressing a membrane transporter NTCP. Biochemical and Biophysical Research Communications, 2014. 443(3): p. 808–813.

7. Cagno, V., et al., Inhibition of herpes simplex-1 virus replication by 25-hydroxycholesterol and 27-hydroxycholesterol. Redox Biology, 2017. 12: p. 522–527.

8. Civra, A., et al., Inhibition of pathogenic non-enveloped viruses by 25-hydroxycholesterol and 27-hydroxycholesterol. Scientific Reports, 2014. 4(1).

9. Shawli, G., et al., The Oxysterol 25-Hydroxycholesterol Inhibits Replication of Murine Norovirus. Viruses, 2019. 11(2).

10. Civra, A., et al., 25-Hydroxycholesterol and 27-hydroxycholesterol inhibit human rotavirus infection by sequestering viral particles into late endosomes. Redox Biol, 2018. 19: p. 318–330.

11. Li, C., et al., 25-Hydroxycholesterol Protects Host against Zika Virus Infection and Its Associated Microcephaly in a Mouse Model. Immunity, 2017. 46(3): p. 446–456.

12. Montgomery, S.R., et al., A novel osteogenic oxysterol compound for therapeutic development to promote bone growth: activation of hedgehog signaling and osteogenesis through smoothened binding. J Bone Miner Res, 2014. 29(8): p. 1872–85.

13. Scott, T.P., et al., Comparison of a novel oxysterol molecule and rhBMP2 fusion rates in a rabbit posterolateral lumbar spine model. Spine J, 2015. 15(4): p. 733–42.

14. Buser, Z., et al., Effect of Oxy133, an osteogenic oxysterol, on new bone formation in rat two-level posterolateral fusion model. Eur Spine J, 2017. 26(11): p. 2763–2772.

15. Wang, F., F. Stappenbeck, and F. Parhami, Inhibition of Hedgehog Signaling in Fibroblasts, Pancreatic, and Lung Tumor Cells by Oxy186, an Oxysterol Analogue with Drug-Like Properties. Cells, 2019. 8(5).

16. Stappenbeck, F., et al., Inhibition of Non-Small Cell Lung Cancer Cells by Oxy210, an Oxysterol-Derivative that Antagonizes TGFβ and Hedgehog Signaling. Cells, 2019. 8(10).

17. Du Toit, A., Coronavirus replication factories. Nat Rev Microbiol, 2020. 18(8): p. 411.

18. Wolff, G., et al., A molecular pore spans the double membrane of the coronavirus replication organelle. Science, 2020. 369(6509): p. 1395–1398.

19. Blanchard, E. and P. Roingeard, Virus-induced double-membrane vesicles. Cell Microbiol, 2015. 17(1): p. 45–50.

20. Matsuyama, S., et al., Enhanced isolation of SARS-CoV-2 by TMPRSS2-expressing cells. Proceedings of the National Academy of Sciences, 2020. 117(13): p. 7001–7003.

21. Marcello, A., et al., The cholesterol metabolite 27-hydroxycholesterol inhibits SARS-CoV-2 and is markedly decreased in COVID-19 patients. Redox Biol, 2020. 36: p. 101682.

22. Arca, M., et al., Increased plasma levels of oxysterols, in vivo markers of oxidative stress, in patients with familial combined hyperlipidemia: reduction during atorvastatin and fenofibrate therapy. Free Radic Biol Med, 2007. 42(5): p. 698–705.

23. Akpovwa, H., Chloroquine could be used for the treatment of filoviral infections and other viral infections that emerge or emerged from viruses requiring an acidic pH for infectivity. Cell Biochemistry and Function, 2016. 34(4): p. 191–196.

24. Liu, J., et al., Hydroxychloroquine, a less toxic derivative of chloroquine, is effective in inhibiting SARS-CoV-2 infection in vitro. Cell Discov, 2020. 6: p. 16.

25. Kokic, G., et al., Mechanism of SARS-CoV-2 polymerase stalling by remdesivir. Nature Communications, 2021. 12(1).

26. Paul, D. and R. Bartenschlager, Architecture and biogenesis of plus-strand RNA virus replication factories. World J Virol, 2013. 2(2): p. 32–48.

27. Zhang, Z. and S. Urban, New Insights into Hepatitis D Virus Persistence: the Role of Interferon Response and Implications for Upcoming Novel Therapies. J Hepatol, 2020.

28. Ogando, N.S., et al., SARS-coronavirus-2 replication in Vero E6 cells: replication kinetics, rapid adaptation and cytopathology. J Gen Virol, 2020. 101(9): p. 925–940.

29. Nao, N., et al., Consensus and variations in cell line specificity among human metapneumovirus strains. PLoS One, 2019. 14(4): p. e0215822.

30. Goto, K., et al., Evaluation of the anti-hepatitis C virus effects of cyclophilin inhibitors, cyclosporin A, and NIM811. Biochem Biophys Res Commun, 2006. 343(3): p. 879–84.

31. Corman, V.M., et al., Detection of 2019 novel coronavirus (2019-nCoV) by real-time RT-PCR. Euro Surveill, 2020. 25(3).

32. Mizutani, T., et al., Phosphorylation of p38 MAPK and its downstream targets in SARS coronavirus-infected cells. Biochem Biophys Res Commun, 2004. 319(4): p. 1228–34.

33. Ohashi, H., et al., The aryl hydrocarbon receptor-cytochrome P450 1A1 pathway controls lipid accumulation and enhances the permissiveness for hepatitis C virus assembly. J Biol Chem, 2018. 293(51): p. 19559–19571.

34. Kaneko, M., et al., A Novel Tricyclic Polyketide, Vanitaracin A, Specifically Inhibits the Entry of Hepatitis B and D Viruses by Targeting Sodium Taurocholate Cotransporting Polypeptide. Journal of Virology, 2015. 89(23): p. 11945–11953.

