## Supplementary Information for "Identification of anti-severe acute respiratory syndrome-related coronavirus 2 (SARS-CoV-2) oxysterol derivatives in vitro"

Received: date; Accepted: date; Published: date

**Supplementary Figure S1**

**Supplementary Table S1**

**Supplementary Figure S1.**

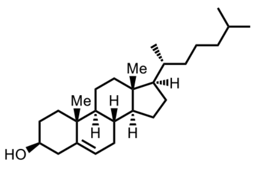

**Cholesterol**

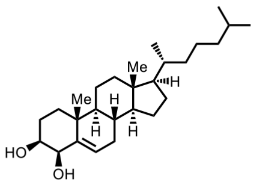

**4beta-OHC**

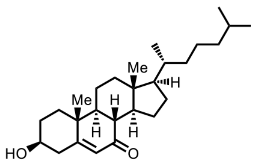

**7-KC**

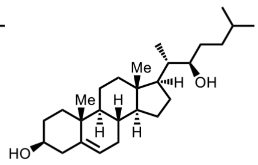

**22(*R*)-OHC**

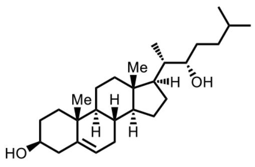

**22(*S*)-OHC**

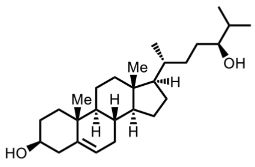

**24(*S*)-OHC**

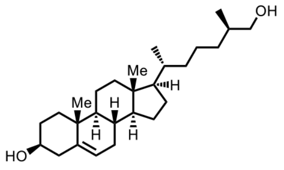

**27-OHC**

**Supplementary Figure S1.**

Chemical structure of the natural oxysterols used in this study.

**Supplementary Table S1**.

**(A) Plasma concentrations of Oxy210**

| Time (h) | Sample concentration (ng/mL) | | | | | Mean ± SD |
|---|---|---|---|---|---|---|
| 24 | 26 | 267 | 8 | 12 | 83 | 79 ± 109 |
| 48 | 1039 | 652 | 1218 | 764 | 622 | 859 ± 259 |
| 96 | 1157 | 771 | 4471 | 6050 | 959 | 2682 ± 2423 |

**(B) Liver concentrations of Oxy210**

| Time (h) | Sample concentration (ng/g) | | | | | Mean ± SD |
|---|---|---|---|---|---|---|
| 96 | 5227 | 3470 | 7302 | 13291 | 5055 | 6869 ±1717 |

**(C) Lung concentrations of Oxy210**

| Time (h) | Sample concentration (ng/g) | | | | | Mean ± SD |
|---|---|---|---|---|---|---|
| 96 | 1688 | 1109 | 6083 | 10524 | 1282 | 4137 ± 1843 |

**Supplementary Table S1**. **Concentration of Oxy210 in plasma, liver, and lung in mice.** Oxy210 mixed in regular Chow food at 4 mg/g of chow was fed to male C57BL/6 mice ad libitum. (**A**) Blood was drawn after 24 h, 48 h and 96h to analyze the plasma concentration of Oxy210. After 96 h, terminal liver (**B**) and lung (**C**) tissue samples were analyzed for Oxy210 concentrations. The data show the concentrations of Oxy210 in each tissue in five mice and its mean values and SD.
